# A Thermodynamic Interpretation of the Stimulated Raman Spectroscopic Signature of an Action Potential in a Single Neuron

**DOI:** 10.1101/2020.04.20.052332

**Authors:** Shamit Shrivastava, Hyeon Jeong Lee, Ji-Xin Cheng

**Affiliations:** Department of Engineering Science, University of Oxford, Oxford, UK; Institute for Sound and Light, Rosalind Franklin Institute, Harwell, UK; Department of Biomedical Engineering, Boston University, MA USA; Department of Electrical and Computer Engineering, Boston University, MA USA

## Abstract

It has previously been suggested that the plasma membrane condenses and melts reversibly during an action potential in a neuron, analogous to an acoustic wave travelling in the compressive membrane region. If true it has fundamental consequences for our understanding of the regulation of biological functions during an action potential. It has long been known that the electrical dipoles in the neuronal membrane reorient during an action potential, observed through a variety of optical methods. However, this information has been insufficient to confirm if and how the collective thermodynamic state of the neuronal membrane changes during an action potential. Here, we show that hyperspectral stimulated Raman spectroscopy (SRS) can resolve the thermodynamic state of the neuronal membranes in a single neuron during an action potential. These measurements indicate that the system becomes ordered and compressed during the de-polarisation phase and disordered and expanded during hyper polarisation Therefore, the observation is consistent with the acoustic hypothesis and SRS provides a powerful tool to not only further validate the hypothesis in future, but also explore the role of membrane thermodynamics during an action potential.

## Introduction

Exciting a neuron causes all – or – none voltage spikes that can be measured between two electrodes that are placed across the neuronal membrane. These voltage spikes are known as action potentials and are generated in the neuronal membrane(1). The present understanding of the phenomenon of action potentials originated from extensive empirical studies on the electrochemical properties of nerves(2). However, these spikes can also be measured using nonelectrical methods such as changes in displacements(1, 3), turbidity, birefringence(4), fluorescence(1, 5), magnetic field(6), force(1), as well as temperature(1, 7). While these nonelectrical aspects of the action potential are well known, they are believed to be non – essential for the biological functions of the action potential. Moreover, the observations themselves are explained as epiphenomenon driven by the voltage spikes using empirical coupling coefficients(8). The model allows for mechanical wave-front to follow the ionic wave front, however a compression wave is the fastest displacement wave that can travel in a system as determined by the speed of sound (9). From a material perspective, changes in temperature and displacement are fundamental to the understanding of any spike or wave propagation phenomenon, as required by the conservation laws of mass, momentum, and energy(10). For example, the present understanding describes the action potential as a purely dissipative phenomenon in analogy to the burning of a fuse where the wavefront propagates as a result of irreversible and diffusive mass (ion) transfer. Hodgkin described in 1964, “The invariance of the action potential arises because the energy used in propagation does not come from stimulus but is released by the nerve fibre along its length. In this respect nervous conduction resembles the burning of a fuse of gunpowder and is unlike current in an electrical cable”(11). Disregarding the “energy of the stimulus” is equivalent to disregarding the role of reversible momentum transfer as seen, for instance, during a sound wave propagation. However, heat studies in nerves indicate that an action potentials is a substantially reversible process(7, 12). In fact, Lichtervelde et al.(13) recently showed that the magnitude of heats can be explained entirely if the process is assumed to be *completely reversible* and the redistribution of ions in the solvation layer is also accounted for. Such a description will be consistent with the coupling of the membrane’s ionic environment with a propagating acoustic wave as previously observed in a lipid monolayer(14). However, to address the question of reversibility it is important to address the efficiency of the process, or the ratio of irreversible to reversible heat. (12). The observed ratio of irreversible:reversible heat in nerves, is of the order of 1:10. Later measurements have observed this value to be even smaller at 6:230(15). Whereas the efficiency calculated on the basis of Hodgkin and Huxley models is of the order of 1:1, i.e. one order magnitude difference. Note that the calculation includes both the contribution from capacitive charging and entropic contribution of conformational changes in ionophores. Therefore, the electrochemical understanding of action potentials that ignores the role of heat and momentum transfer during the action potential has long been debated(1, 16–18).

Recent research has brought the thermodynamic underpinnings of the phenomenon at the centre of this debate by describing the nerve impulse as a material wave that propagates as a localized condensation of the membrane(19, 20). The theory incorporates all the non-electrical aspects of the nerve impulse as default thermodynamic couplings and explains the observed reversible heating and cooling as a default consequence of adiabatic compression and rarefaction of the medium during pulse propagation. A particular prediction of the theory is a change in the state of the matter so significant that the membrane’s thermodynamic susceptibilities, such as heat capacity and compressibility, change significantly. Such significant changes in the compressibility are observed during thermotropic transitions in biological(21, 22) as well as artificial lipid interfaces(23). Under such conditions the nonlinear properties of the action potentials, such as all – or – none excitation and annihilation of pulses upon collision, are also shown by compression waves in artificial lipid films(24, 25), indicating that the “invariance of action potentials” can arise also from momentum transfer. Therefore, the theory predicts a condensation (freezing) of the membrane during an action potential as it depolarises and a rarefaction (melting) as it polarises again. The hyperpolarisation is then a consequence of the inertia as the membrane relaxes(24, 26). However, a direct measurement of the membrane state to confirm its freezing during an action potential has remained elusive. Here, by observing the changes in a signature Raman peak of the plasma membrane, we confirm that the thermodynamic state of the membranes changes significantly, during an action potential in a single neuron.i.e. the collective order of the membrane undergoes cyclic change in a manner that is consistent with the acoustic hypothesis.

## Measuring thermodynamic state changes from light-matter interaction

Optical methods have a long history in measuring physical changes in the membrane during an action potential(5, 27). The electromagnetic nature of light allows probing the changes in the electric field around the dipoles, either extrinsic or intrinsic to the membrane, due to changes in membrane potential. Thus changes in optical signals, in general, can be easily mapped to changes in membrane potentials, which form the basis for many important methods for studying action potentials. However, such light-matter interactions do not lie outside the purview of thermodynamics(28, 29), and the properties of light can indeed be used as thermodynamic observables of the system that they interact with(30, 31).

When photons interact with a material, a small fraction of their population changes its wavelength due to the second-order effects of the thermal fluctuations in the material(32). These perturbed photons lie on a spectrum and the frequency distribution of the photons directly represents the partition of thermal energy among the conformational states. Raman and infrared spectroscopy-based methods are well known to characterize the conformational state of artificial and biological membranes (33–35). Here, we exploit this relationship to obtain new insights into the nature of physical changes in neuronal membranes during an action potential.

We start with fact that the entropy of a membrane complex can be written as a thermodynamic function of the form *S* ≡ *S*(*x_i_*)(31, 36), where *x_i_’s* are the extensive observables of the system that can be measured experimentally, e.g. volume, number of charges, number of photons, enthalpy, and energy of compartments etc.

Using this the Taylor expansion of the entropy potential can be written as:

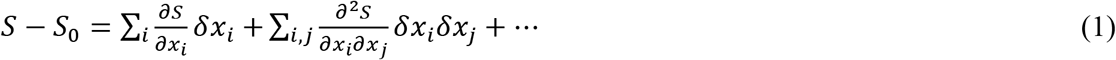

We employ the equation under the assumption of local equilibrium where the state variables are defined spatio-temporally and 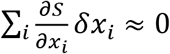 (37, 38). Then using eq. (1) and the inversion of Boltzmann principle *w* = *e^S/k^* it can be shown that various observables must be coupled by the relation

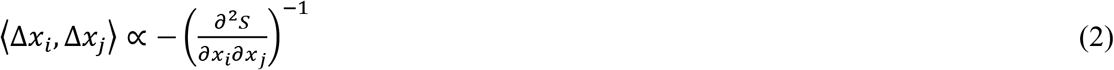

The approach previously explained the changes in the emission spectra of solvato-chromic dyes embedded lipid vesicles exposed to shock waves(31). In this case the entropy potential was described as a function of the emission spectra of the dye, representing the distribution of the ground state dye molecules in equilibrium with the system.

So how do these equations help us in understanding thermodynamic changes in membranes using Raman spectroscopy? If the Raman spectrum is represented by the function *I_n_*, i.e. the number of photons at wavenumber *n*, then as per the above, we propose the ansatz that the entropy of a complex membrane can also be written as *S* = *S*(*I_n_*, *x_i_*) (28, 39). Since all the arguments of the function are experimentally measurable, the function provides a general basis for designing experiments, where *x_i_* are chosen depending on the phenomenon of interest. For example, *I_n_* can be measured as a function of temperature, *T*, and in that case eq.(2) gives;

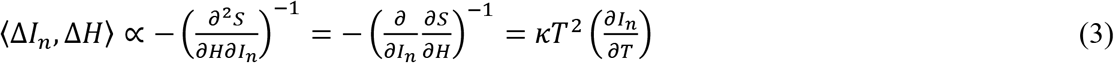

i.e. small changes in intensity at a given wavenumber Δ*I_n_* for a given change in the total enthalpy of the system, Δ*H* are required to obey eq.(3) as per the second law of thermodynamics. Here *κ* is the Boltzmann constant and 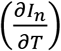 represents the derivative of the experimentally measured function *I_n_*(*T*). Therefore, for conformational changes that occur during a process in the membrane, as observed from Δ*I_n_*, an equivalent enthalpy change can be estimated from Eq. (3). Similarly, *I_n_* can also be investigated as a function of pressure or electric field to reflect changes in volume or membrane potential.

## State changes during voltage-clamped experiments

A voltage-clamp experiment can be assumed to be a constant temperature or isothermal experiment (at the temperature of the sample chamber), while other variables are free to adjust (e.g. membrane pressure or interfacial pH). Then the changes in state as per eq. (2) are due to changes in the electrical field applied to the membrane. Recently, Lee and Cheng reported spatially and temporally resolved *I_n_* measurements on single neurons using stimulated Raman scattering (SRS) spectroscopy(40). As discussed above, the data was interpreted by studying *I_n_* as a function of membrane potential, showing that structural changes in a single neuron can be resolved during an action potential based on observed Δ*I_n_*. Figure 1c shows a typical SRS spectrum of a neuron at the resting potential, *I_n_* for *n* ∈ [2800,3000]*cm*^−1^ indicating the strongest contribution from a peak at 2930 *cm*^−1^. Figure 1e shows the changes in the SRS spectra Δ*I_n_* measured during somatic voltage clamp of a neuron (−80 *mV to* + 30 *mV*) with respect to the resting potential of −60*mV*. Thus, the intensity of 2930 *cm*^−1^ peak decreases in response to membrane depolarisation and increase upon hyperpolarisation.

**Figure 1.**
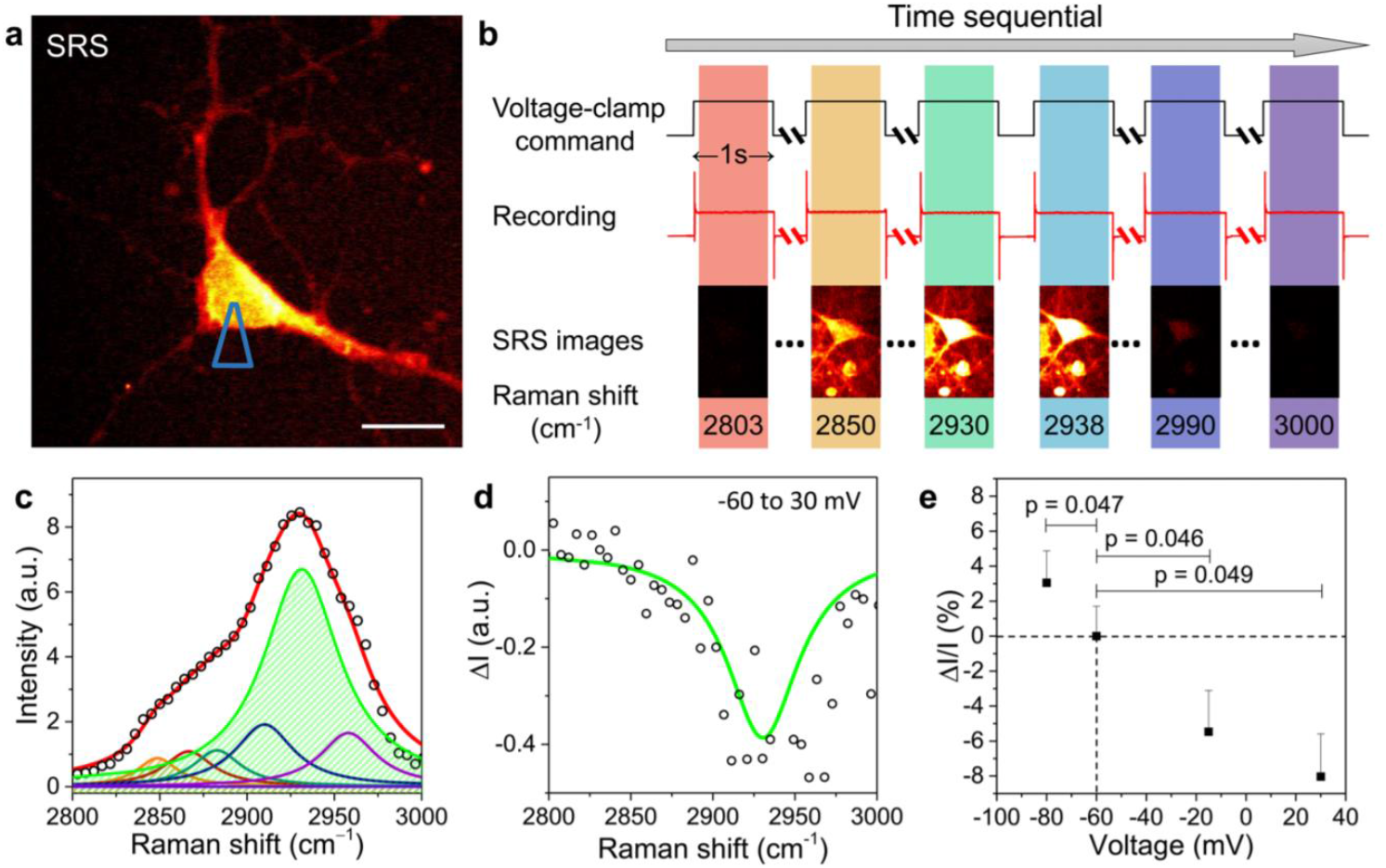
(a) SRS image of a patched neuron with micropipette position indicated. Scale bar: 10 μm. (b) Schematic of hyperspectral SRS imaging of neurons while holding different potentials. (c) Representative SRS spectrum of a patched neuron (dot), fitted using seven Lorentzian (colored lines) with major contributing bands filled, 2850 (orange) and 2930 cm^−1^ (green). Red: cumulative fitted curve. (d) SRS spectral change, ΔI (dots), of the neuron from −60 to +30 mV with fitted curve (line). (e) Percentage changes of SRS intensity (ΔI/I) of neurons at 2930 cm^−1^ at various membrane potentials. Error bars: + standard error of the mean (SEM). Reprinted (adapted) with permission from (J. Phys. Chem. Lett. 2017, 8, 9, 1932-1936). Copyright (2017) American Chemical Society

A potential mechanism was previously provided in terms of interaction of the Fermi-resonance component of the 2930 *cm*^−1^ peak with the changes in the ionic environment during depolarisation. However, the magnitude of the effect was significantly less than changes observed during the voltage clamp experiment. Here, we provide a mesoscopic, empirical, and thermodynamic interpretation of these changes based on eq. (3). A salient feature of the approach as discussed above is that many effects of microscopic origin, which might all possibly contribute to observed changes in the Raman spectra, are implicitly accounted for the macroscopic entropy potential, including the changes in ionic environment, polarity, or any other microscopic agents of change. The analytical and logical consistency exists between the empirical equations that are independent of any molecular models. For example, eq. (3) does not assume the underlying molecular structure of the membrane, i.e. it is invariant of the molecular origin of Raman signal which could be due to any protein, lipids, ion, etc. This invariance provides us with means to refer to model systems for simulating Δ*I_n_* (as observed in real neurons) and derive new insights from it. In this sense, changes in *I*_2930_ as a function of the state has been measured in a variety of artificial as well as biological membranes. Both in lipids and proteins, the band represents a “melting marker” as the peak at 2930 *cm*^−1^ appears in difference spectra Δ*I_n_* only when the corresponding state change involves a cooperative thermotropic transition or melting(41). These studies(42–45) usually measure intensity around 2930 *cm*^−1^ relative to intensity at another wavenumber, which provides an internal reference that removes systematic errors. Note that thermotropic transitions are not exclusive to pure lipid membranes or proteins, the cooperativity during the transitions is long known to extend over domains or proto-mers consisting of both proteins and lipids(42, 46). Furthermore, Δ*I_n_* have been measured for thermotropic transitions in membranes and proteins induced by a variety of thermodynamic fields including temperature(41, 44, 47), pressure(45), as well as pH(42). Thus, the term “thermotropic transition” is employed in the most general sense, which is indicated by a nonlinearity or cooperativity in the functional relationship between the thermodynamic variables. For example, by observing nonlinearities in *I*_2930_ as a function of pH and temperature in erythrocytes ghosts, others have previously concluded that Δ*I*_2930_ represents a “concerted process at apolar protein-lipid boundaries”(42, 43). Thus a change in *2930 cm*^−1^ peak can represent a change in the order of either lipids, proteins, or their combination.

Therefore, Δ*I*_2930_ in figure 1 shows that the membrane essentially undergoes a collective increase in order during voltage-clamped depolarisation. That is, just like temperature, pressure, and pH; membrane potential can also induce an ordering effect in the membrane. Furthermore, based on observed Δ*I*_2930_ in figure 1 the extent of ordering can also be estimated. For example, (*I*_2930_/*I*_2850_) changes from 1.9 at −60*mV* (resting potential) to 1.6 at +30*mV* (depolarised), i.e. ~18% change. Temperature dependence of Raman spectra has been measured previously in a variety of neuronal membranes(48, 49). For example, in unmyelinated garfish olfactory nerve, the ratio (*I*_2950_/*I*_2885_) was measured as a function of temperature. The ratio changes from 1.22 at 25 *°C* to 1.09 at 6 *°C*, i.e. 12% change over ΔT = 19 °*C*, which as discussed in detail by the authors, represents a significant increase in order. On the other hand, based on the SRS study in primary neurons by Lee et al.(40), (*I*_2950_/*I*_2885_) changes from 1.18 at −60*mV* (resting potential) to 1.04 at +30*mV* (depolarised), i.e. 13% change.. It is important that unlike other optical observables measured previously, where the sign of the signal could not be interpreted in absolute terms, a negative Δ*I*_2930_ has an absolute meaning which indicates an increase in order.

Using these numbers, we estimate the enthalpy of phase change as well as average heat capacity of neuronal membranes from *I_n_*(*T*) using the Van’t Hoff’s approximation (50) and equilibrium constant defined in terms of Raman intensities (51);

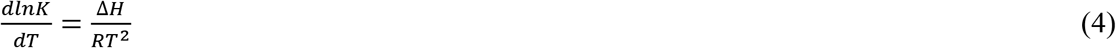

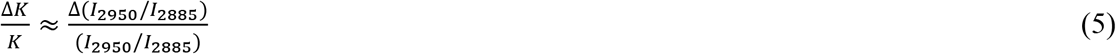

A 12% change in (*I*_2950_/*I*_2885_) over Δ*T* = 19 *°C* gives an average Δ*H* ≈ 4.5 *kJ/mol* and *c_P_* = 224*Jmol*^−1^*K*^−1^. These values are now employed to estimate state changes during an action potential.

## State changes during an action potential

Can the analysis be extended to state changes during an action potential? Raman spectroscopy has a long history of determining dynamic state changes within propagating wavefronts in fields like thermos-fluids(52–54). While the above discussion assumed equilibrium during voltage clamp, eq. (2) can be extended to an arbitrary macroscopic state (partial equilibrium)(38); by observing region so small that the corresponding relaxation time (≈ *l/c, l* is the length of the region, and *c* is the speed of sound in the medium) is smaller than the fastest timescale of interest in the underlying process. SRS signal (fig. 2) obtained previously during an action potential in a single neuron satisfies these criteria and showed Δ*I_2930_ ≈* −1% upon depolarisation and Δ*I*_2930_ ≈ +1% upon hyperpolarisation during an action potential.

**Figure 2.**
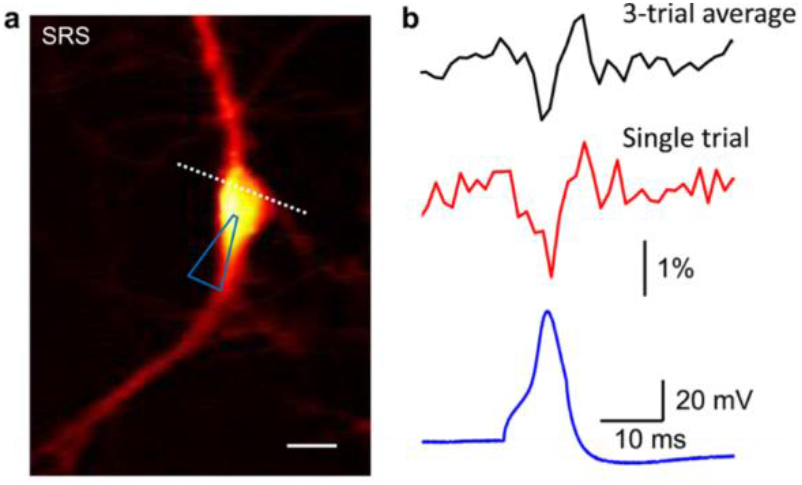
(a) SRS image of a patched neuron. The dashed line indicates the scanning trace. (b) 3-trial average (black) and single-trial SRS time trace (red) of the neuron shown in panel a with simultaneous current clamp recording (blue), showing a single action potential. The SRS intensity was normalized by the SRS reference generated with the same pulses. Scale bar: 20 μm. Reprinted (adapted) with permission from (J. Phys. Chem. Lett. 2017, 8, 9, 1932-1936). Copyright (2017) American Chemical Society

Note that unlike the voltage-clamp experiments where the temperature is constant, there is no such requirement during an action potential, which represents a fundamentally different thermodynamic process. To compare the two, recall that there are quantities that can remain constant even during such dynamic process, most well-known being the entropy that remains constant, for example, during the propagation of sound waves. While an infinitesimal acoustic perturbation is completely reversible, finite amplitude sound waves, for example shock waves, are dissipative. However, even in the case of shock waves the entropy production depend on third order changes in volume and pressure. As a result, state change across a shock front is still adiabatic to second order. (55, 56). Furthermore, the observed temperature changes during a nerve impulse indicate a substantially reversible phenomenon, which led to the original suggestion that nerve impulse or action potentials emerge from the same underlying physics as sound waves(16).

## Relation to the thermodynamics of compression waves and fluctuations

So if the state changes during an action potential are indeed similar to those that might occur during sound waves, how do we reconcile (i) the membrane freezes, while (ii) the temperature increases, (iii) the entropy remains constant, and (iv) channel activity increases? Critical insights on this matter have been generated by studying 2D compression waves and fluctuations in artificial lipids films.

Let us first consider the compression waves. Remarkably, near an order-disorder transition in a lipid film, these waves show many key characteristic properties of action potentials including all-or-none excitation(20, 24) and annihilation upon collision(25). In the case of artificial lipid films, the characteristics emerge naturally from the underlying thermodynamic state changes. The picture that has emerged is plotted in figure 3. Note how the phase transition region can be traversed differently at a constant temperature, constant entropy, or constant pressure. Consider the initial state of the membrane represented by (*T*_0_*S*_0_), then Δ*I*_2930_ ≈ −9% during the voltage clamp represents condensation along a constant temperature path (*T* = *T*_0_), while Δ*I*_2930_ ≈ −1% during the action potential represents condensation along constant entropy path (*S* = *S*_0_). It is important to recall that conventionally the membrane state are velieved to be the same at the same voltage during a voltage clamp experiment and during an action potential, however this is inconsistent with the above changes in Raman spectra.

**Figure 3.**
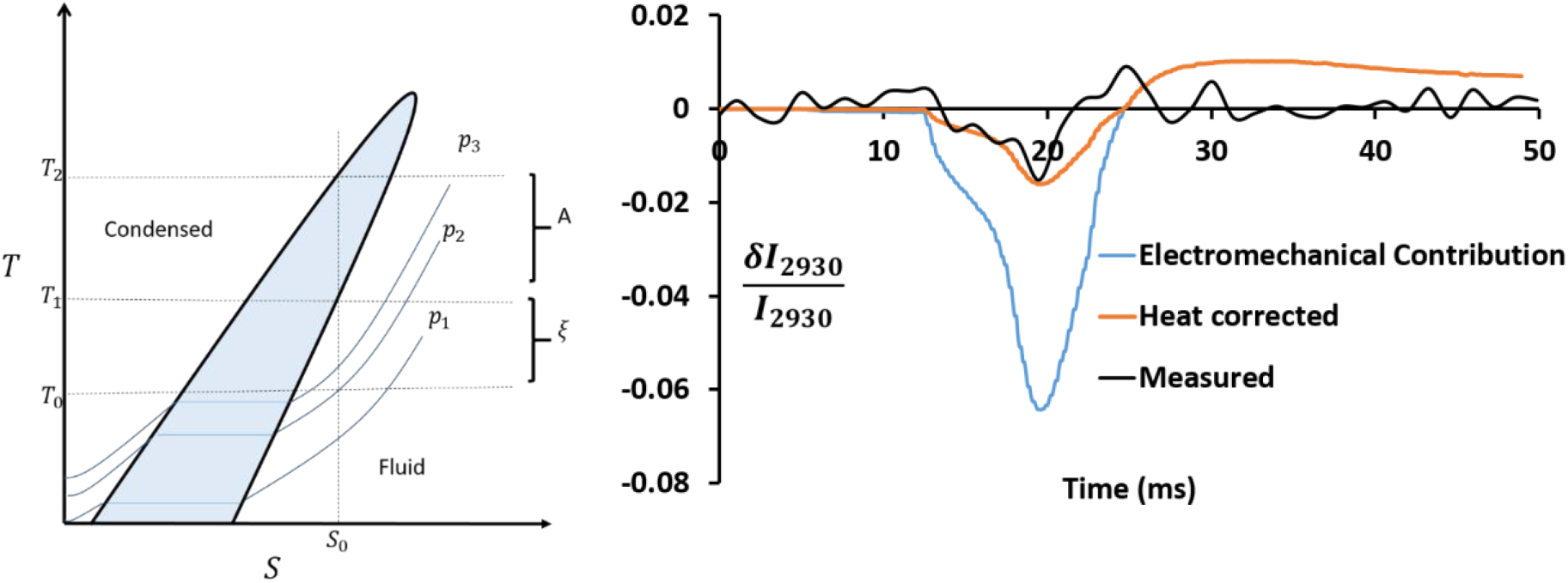
(Left) TS diagram as an aid to the eye, informed by experiments in artificial lipid films*(9, 20)*. The hypothesized transition region is shaded. The region makes an acute angle with S axis for retrograde materials. Assuming that complete phase change takes place across the wavefront where the system is always in local equilibrium during an impulse, starting at (T_*0*_S_*0*_), the process across the wavefront is represented by the isentrope S = S_*0*_. The threshold and amplitude scale with ξ and A respectively, i.e. the distance between the initial state and the intersection of the isentrope with the phase boundary. Different isobars p_i_ have been overlaid where p_*3*_ > p_*2*_ > p_*1*_. (Right) Purely electrochemical component δI_*2930V*_(t) (blue curve) estimated from voltage clamp experiments (from fig.1), is overlaid on calculated heat corrected curve δI_*2930*_ (t)_heat corrected_, and measured δI_*2930*_ (t) (from fig.2)

Therefore, the competition between the condensing effect of the electromechanical compression of the membrane and the fluidizing effect of the increase in temperature due to the release of latent heat is determined by the inclination of the co-existence region in the TS diagram, a parameter known as retrogradicity(20, 57). Another way to interpret the behaviour is in terms of heat capacity; if the heat capacity of a material is sufficiently high, the temperature rise due to the latent heat (released during condensation) will be small enough for the system to still condense under compression. For example, water vapour does not condense upon adiabatic compression. On the other hand, polymers with chain lengths greater than 4 carbon atoms, typically show retrograde behaviour and condense upon adiabatic compression (*c_P_* > ~200*Jmol*^−1^*K*^−1^).(58)

Therefore, the observed *δI*_2930_ (*t*) has thermal as well as electromechanical contributions that can be estimated to first order using;

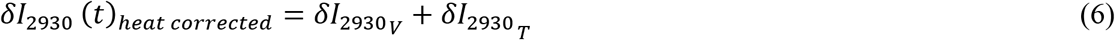

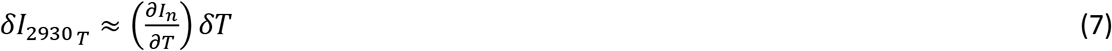

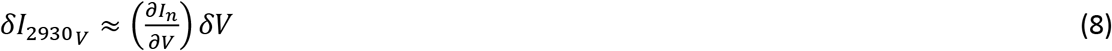

To evaluate these equation we make following assumptions; (i) the resting state is close to the phase boundary, (ii) due to latent heat contributions Δ*H* is assumed large compared to state changes that do not involve a phase change, (iii) *δV* (*t*) is assumed to span the phase change region completely, i.e. the fraction of membrane in condensed phase, *α* = 0 at *δV*(*t*) = −60*mV* and *α* = 1 at *δV*(*t*) = 30*mV*, (iv) 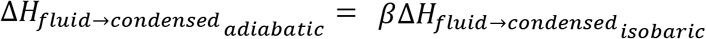, where is *β* is a phenomenological parameter <1 as adiabatic phase change is closer to the apex of the coexistence dome (enthalpy of transition decreases as the transition temperature approaches critical temperature) and we assume 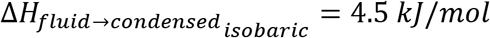 i.e. the estimate obtained above. With these assumptions we get;

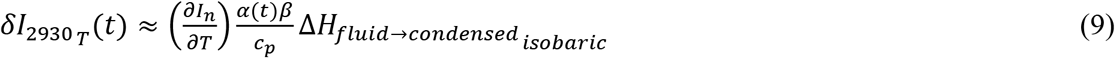

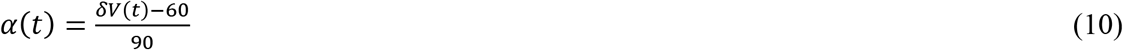

*δI*_2930_ (*t*)*_heat corrected_* thus calculated is plotted in figure3 for *β* = 0.4. As shown the measured *δI*_2930*τ*_(*t*) is recovered from the voltage clamp data after accounting for the latent heat during an adiabatic phase change. It is interesting to note that during the rising phase the estimated and observed *δI*_2930_(*t*) align correctly, the recovery in measured *δI*_2930_(*t*) occurs before the recovery of *δI*_2930_V__(*t*) indicating that membrane relaxation occurs before the recovery in voltage.

That the orientation of dipoles in the membrane changes was long known through an extensive range of studies, however a change in dipole orientation is necessary but not sufficient to conclude an increase in order(30, 59). For that, changes in both the magnitude and the distribution of an observable are required as indicated by eq. (2). Few previous studies have performed such experiments on nerves however a corresponding thermodynamic interpretation was not provided. For example, measurements of difference in infrared spectra between resting and excited state of nerve bundles from several species indicated changes in the conformation of membrane phospholipids(60). Similarly, Raman peaks of carotenoids(61) in the membrane during an action potential indicated compression of the molecules, however, whether the compression corresponds to freezing could not be inferred. Tasaki et.al. measured the difference in the emission spectrum between the resting and excited state of solvatochromic fluorescent dyes(62). While the observed blue shift indicated an increase in order, the recent thermodynamic interpretation of the spectral width of these dyes(39) confirmed that an increase in order was indeed observed in Tasaki’s experiments. However, given the uncertainty regarding the specific environment of dyes, doubts remained. Similarly, an order-disorder transition during an action potential has been indicated by the measurements of membrane-fluidity sensitive dyes(63).

Finally, a brief comment on the nature of fluctuations during the process which relates to channel activity during an action potential. Here, Δ*I*_2930_ represents first-order changes in the state during an action potential. However, fluctuations represent second-order changes, which have been proposed as the basis for the regulation of channels current (64–67) in the thermodynamic approach and can be accessed by measuring the second order changes ΔΔ*I*_2930_, e.g. by resolving changes in the width of the peak(31). The relation between fluctuation and temperature includes a material constant (see eq. (2) and eq. (3)), like heat capacity or compressibility (in general known as thermodynamic susceptibilities). Usually, susceptibilities are taken to be “constants” and fluctuations are assumed to be a function of temperature alone. However, the assumption breaks down near a phase transition where heat capacity and compressibility of the membrane have large peaks(21). As the membrane freezes, it goes through this intermediate peak in susceptibility (metastable states), which will increase fluctuations that should cause a spike in channel currents. Such transitions have also been referred as pseudocritical transitions in lipid membranes. Direct evidence of such a role of channel currents during an action potential remains to be seen.

## Challenges and Limitations

Here we have provided a thermodynamic interpretation of the changes in the phenomenological Raman intensities as a function of the state of the entire system.However, unveiling the material basis of these changes beyond the abstract lipid-protein protomers as hypothesised by Changeux et.al. (46) is challenging. The observed region of the spectrum (fig.1c) is dominated by the protein band at 2930*cm*^−1^, which obfuscates information specific to lipid band at 2850*cm*^−1^.(68) The interpretation in terms of thermotropic transition is mainly based on the Fermi resonance component of the chain terminal methyl C-H symmetric stretching mode. This is a known challenge that will requires characterising other regions, in particular, 1000 to 1200*cm*^−1^ region related to C-C skeletal stretching. However, this band may present other challenges, for example, the presence of strong resonance Raman bands of the carotenoid pigment.(48) Furthermore, the observed protein signal can be from both the membrane and the axo-plasm that lie within the optical focal spot. Here, it has been assumed based on the timescales, that the changes in the spectra during an action potential arise mainly from components in the plasma membrane that are directly involved in the excitation process. This assumption is consistent with the source of optical changes in nerve fibres, in general, which have been shown to lie within or close to the surface of the membrane and not in the axo-plasm(69).

The quantitative assumptions made in above interpretation also underline the limitations of this study. The co-efficient 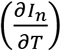 was estimated based on measurements in rat sciatic nerve and unmyelinated garfish olfactory nerve, assuming that it holds generally(49). However, ideally it needs to be measured within the same sample. Furthermore, we assumed that changing the temperature and the voltage of the clamp affect the same system as propagating action potential, and that local equilibrium can be assumed. These assumptions can been justified considering the objective was to estimate orders of magnitude of the enthalpy changes, which were tested against the condition 0 < *β* < 1. Also the same assumption is implicit when extrapolating coefficients of voltage clamp measurements to propagating action potentials in electrophysiology.

The nature of the observed thermotropic transition also needs to be firmly established to understand its material basis. Observing several other region of the spectra over a wider range of temperature and voltages, as well as other variables, such as pH and pressure, will be required to reveal the nonlinearities associated with the claimed thermotropic transition. Furthermore, a definite proof of phase transition during the action potential based on Raman spectra would also require observing time resolved changes in the shape of the spectra. Direct measurement of the changes in the width of the spectra can confirm the extent of phase change during the action potential. However, considering that even in pure lipid membrane, typical shifts in the spectra are of the order of 1*cm*^−1^, its presents a formidable technological challenge. There are other fundamental aspects of the phase transition as well that require further investigations. For example, phase boundaries appear as single points in single component systems when observed as a function of a single variable, but in multicomponent system phase-boundaries exist as hypersurfaces and it is important to approach these boundaries using multiple variables for a complete understanding(70–72). For example, using solvato-chromic dyes it was recently show that in Chara Alga, the state of the membrane may change in strongly nonlinear manner as function of pH even though it changes linearly as a function of temperature (73). Relative changes in 2930*cm*^−1^ band, both with respect to the temperature and voltage, have been found to change linearly in neurons over the limited range they have been tested. However, observed change in 2930*cm*^−1^ band are found to be significant based on the corresponding changes observed during transitions in a range of lipids, fatty acids(74), proteins(43), and membrane systems(42).

Finally, going back to the burning fuse analogy of action potential, the material in general expands during such a process where as this study shows a reversible compression (and a thermotropic condensation), making it highly unlikely that wave front propagation does not involve momentum transfer(75). However, answering that question completely will require systematic investigation of the entire phenomenon as a function of thermodynamic state (76). Most important would be to show the dependence of conduction velocity on a thermodynamic susceptibility. The Raman spectra as shown here provides access to a multi-dimensional internal state observable of the system, which will be critical for testing the hypothesis. Thermodynamics lays down strict conditions for supersonic or sonic vs subsonic propagation and thus provide a critical test for acoustic vs ionic hypothesis. While sonic or supersonic propagation requires a compression of the material across the wavefront; a subsonic propagation, as in ionic hypothesis, is only allowed under expansion across the wavefront(10). Therefore, the present observations are consistent with the acoustic hypothesis.

In conclusion, SRS measurements for the first time provide direct access to conformational changes in the membrane of a single neuron during an action potential. Therefore, it is now possible to test various predictions of the acoustic hypothesis regarding the role of the thermodynamic state of the membrane in the action potential phenomenon(1, 12, 16, 77, 78). In this study, we showed that the observations so far are consistent with these predictions, however as the technology improves, direct observation of changes in the width and peak positions during an action potential will allow further validation of the hypothesis. These observations will provide further validation of the cooperativity of the observed changes. While the nature of transition across the wave-front remains to be confirmed, the observations strongly suggest a compression and increase in order across the wavefront. Given that a compressive wave-front can propagate either at sonic or supersonic speeds, the observation supports the acoustic hypothesis(79). The observed changes are consistent with the behaviour of two-dimensional compression waves recently observed in artificial lipid films(24). At least in the case of artificial lipid films, the characteristics emerge naturally from the underlying thermodynamic state changes. Now, SRS provides a powerful technique with the spatio-temporal resolution required to investigate action potentials in single neurons as a thermodynamic phenomenon (1, 16, 19, 24).

